## Supplementary material for "Single molecule real time (SMRT) full length RNA-sequencing reveals novel and distinct mRNA isoforms in human bone marrow cell subpopulations": Suppl Fig S1 to S4

**Contents:**

- Supplemental Experimental Procedures
- Figures S1 to Fig S4
- Supplemental References

**- A separate file contains Tables S1 to S7: "Deslattes Genes Tables S1 to S7 Febr 18 2019.xlsx"**
Table S1: Top five transcript isoform counts for genes with 2 to 16 exons (Fig. 5b)
Table S2: Transcript ID, gene names, detection of peptides by mass spectrometry (Fig. 5b and Table S1).
Table S3: Number of exons, transcript isoform numbers (Trans #) for genes in Fig. 5a.
Table S4: Number of exons, transcript isoform numbers (Trans #) for genes in Fig. 5c.
Table S5: Distribution of exon counts and transcript isoform frequency.
Table S6: Validation of the transcript isoforms
Table S7: MatchAnnot analysis of isoform alignments

**Supplemental Experimental Procedures (Scripts):**

**Transcriptome Alignment and Assembly from Illumina Data.**
The specific qsub command for the total bone marrow (T) alignment was:

```
#!/bin/bash
#$ -cwd
#$ -pe orte 32
#$ -N "th.fr.unstr.tot"
#$ -V
export PATH="/mnt/myenv2/bin:/mydata/cufflinks-2.2.1.Linux_x86_64;/mydata/bowtie2-2.2.5:/mydata/samtools-1.2:$PATH"
/mydata/tophat-2.0.14.Linux_x86_64/tophat2 -p 32 -G /results/Homo_sapiens/UCSC/hg19/Annotation/Genes/genes.gtf \
                      -r200 \
                      --library-type fr-unstranded \
                      -o /trimmed_fastq/transcriptome/ill.bm.total/tophat.2.0.14.fr-unstranded.gtf \
                         /results/Homo_sapiens/UCSC/hg19/Sequence/Bowtie2Index/genome \
                         /trimmed_fastq/trimmed.fasta.total.bm.non.ss/bm.trimmed.reads.1.fa \
                         /trimmed_fastq/trimmed.fasta.total.bm.non.ss/bm.trimmed.reads.2.fa
```

The specific command for cufflinks 2 assembly for total bone marrow (T) was:

```
#!/bin/bash
#$ -cwd
#$ -pe orte 32
#$ -N "cf2.tot.fr.un.gtf"
#$ -V
export PATH="/mnt/myenv2/bin:/mydata/cufflinks-2.2.1.Linux_x86_64;/mydata/bowtie2-2.21:/mydata/samtools-0.1.19:$PATH"
/mydata/cufflinks-2.2.1.Linux_x86_64/cufflinks -b /results/Homo_sapiens/UCSC/hg19/Sequence/WholeGenomeFasta/genome.fa \
  -p 32 \
  -g /results/Homo_sapiens/UCSC/hg19/Annotation/Genes/genes.gtf \
  -L tot.fr.unstranded.gtf \
  -o /trimmed_fastq/transcriptome/ill.bm.total/cufflinks2.2.1.tophat.2.0.14.fr.unstranded.gtf.5.19 \
  /trimmed_fastq/transcriptome/ill.bm.total/tophat.2.0.14.fr-unstranded.gtf/accepted_hits.bam
```

The specific qsub command for the lineage-negative (N) alignment was:

```
#!/bin/bash
#$ -cwd
#$ -pe orte 32
#$ -N "th.neg.gtf"
#$ -V
export PATH="/mnt/myenv2/bin:/mydata/cufflinks-2.2.1.Linux_x86_64;/mydata/bowtie2-2.2.5:/mydata/samtools-1.2:$PATH"
/usr/local/bin/tophat -p 32 -G /results/Homo_sapiens/UCSC/hg19/Annotation/Genes/genes.gtf \
                      --library-type fr-firststrand \
                      -r200 \
                      -o /trimmed_fastq/transcriptome/ill.lin.neg/tophat.2.0.14.fr.firststrand.no.mm.gtf \
                         /results/Homo_sapiens/UCSC/hg19/Sequence/Bowtie2Index/genome \
                         /trimmed_fastq/trimmed.fasta.lin.neg.ss/lin.neg.trimmed.reads.1.fa \
```

```
                              /trimmed_fastq/trimmed.fasta.lin.neg.ss/lin.neg.trimmed.reads.2.fa
```

The specific command for cufflinks 2 assembly for lineage-negative (N) was:

```
#!/bin/bash
#$ -cwd
#$ -pe orte 32
#$ -N "cf2.ill.lin.neg.gtf"
#$ -V
export PATH="/mnt/myenv2/bin:/mydata/cufflinks-2.2.1.Linux_x86_64;/mydata/bowtie2-2.21:/mydata/samtools-0.1.19:$PATH"
/mydata/cufflinks-2.2.1.Linux_x86_64/cufflinks -b /results/Homo_sapiens/UCSC/hg19/Sequence/WholeGenomeFasta/genome.fa \
  -p 32 \
  -g /results/Homo_sapiens/UCSC/hg19/Annotation/Genes/genes.gtf \
  -L ill.lin.neg.nm.th \
  -o /trimmed_fastq/transcriptome/ill.lin.neg/cufflinks2.2.1.no.mm.no.u.gtf \
   /trimmed_fastq/transcriptome/ill.lin.neg/tophat.2.0.14.fr.firststrand.no.mm/accepted_hits.bam
```

**Confirmation of novel transcript isoforms with blast**
To confirm novel transcripts isoforms, two blastable databases were prepared separately for the total bone marrow (total.bm.non.ss) and lineage-negative short read RNA sequences (lin.neg.ss) using the example script below. The full length RNA-seq reads for each gene was split into single sequence files then batch blasted in a script such as this used for *ANXA1*:

```
#!/bin/bash
endings="aa ab ac ad ae af ag ah ai aj ak al am an ao ap aq ar as at au av aw ax ay az
         ba bb bc bd be bf bg bh bi bj bk bl bm bn bo bp bq br bs bt bu bv bw bx by bz
         ca cb cc cd cd cf cg ch ci cj ck cl cm cn co"
#rm ANXA1.lin.neg.coverage.txt
#rm ANXA1.total.coverage.txt
for end in $endings;
do
   echo ANXA1.split."$end"
   blastn -db ../lin.neg.ss/lin.neg.ss -perc_identity 99 -ungapped -num_threads 32  -max_target_seqs 10000 -query
ANXA1.split."$end" -outfmt "6 qseqid qstart qend \
qseq sseq sseqid sstart send" -out ANXA1.split."$end".lin.neg.out
   blastn -db ../../total.bm.non.ss/total.bm.non.ss -perc_identity 99 -ungapped -num_threads 32 -max_target_seqs 10000 -
query ANXA1.split."$end" -outfmt "6 qseqid \
qstart qend qseq sseq sseqid sstart send" -out ANXA1.split."$end".total.out
done
```

Using this output file, a *summarize* script provides the coverage.

```
#!/bin/bash
endings ="aa ab ac ad ae af ag ah ai aj ak al am an ao ap aq ar as at au av
ba bb bc bd be bf bg bh bi bj bk bl bm bn bo bp bq br bs bt bu bv
```

```
ca cb cc cd cd cf cg ch ci cj ck cl cm cn co"
for end in endings
do
echo ANXA1.split."$end"
var=$(grep ">" ANXA1.split."$end"
var2=${var:1}
echo "$var2"
grep "$var2" ANXA1.split."$end".lin.neg.out | cut -c -20 > ANXA1.split."$end".lin.neg.num.txt
sort -n -k 2,3 ANXA1.split."$end".lin.neg.num.txt > ANXA1."$end" > . "$var2".lin.neg.txt
sort -n -k 2,3 ANXA1.split."$end".lin.neg.out > ANXA1."$end". "$var2".lin.neg.fa.txt
cat ANXA1."$end" . "$var2".lin.neg.txt >> ANXA1.lin.neg.coverage.txt
grep "$var2" ANXA1.split."$end".total.out | cut -c -20 >ANXA1.split."$end".total.num.txt
sort -n -k 2,3 ANXA1.split."$end".total.num.txt >ANXA1."$end". "$var2".total.txts
sort -n -k 2,3 ANXA1.split."$end".total.out > ANXA1."$end"."$var2".total.fa.txt
cat ANXA1."$end"."$var2".total.txt >> ANXA1.total.coverage.txt
done
```

*Summarize* finds the matches and prepares the coverage file. Next, a series of *awk* scripts find the gaps in coverage as a result of the blast and limits the hits to 100 bp.
The output of the short reads that match each of the novel isoforms were then searched for the full length reads. All genes followed the same stepwise procedure.
step 1 - Simplify the blast output capturing the ends of the hit, and the length of the overlap.

```
cat ANXA1.total.coverage.txt | awk '{print $1 "\t" $2 "\t" $3 "\t" $3-$2+1}' >ANXA1.total.coverage.maths.txt
```

Ensure the hits are at least 100 bp

```
grep "100" ANXA1.total.coverage.maths.txt > ANXA1.total.100.coverage.maths.txt
```

step 2 - preserving the end of the previous line, this awk script identifies gaps.

```
cat ANXA1.total.100.coverage.maths.txt | awk '{print var "\t" $1 "\t" $2 "\t" $3 "\t" $4 "\t" var-$2;var=$3}' >
ANXA1.total.100.coverage.all.txt
```

step 3 - use the end of the match from the previous line to determine if there is a gap

```
grep "\-" ANXA1.total.coverage.all.txt
```

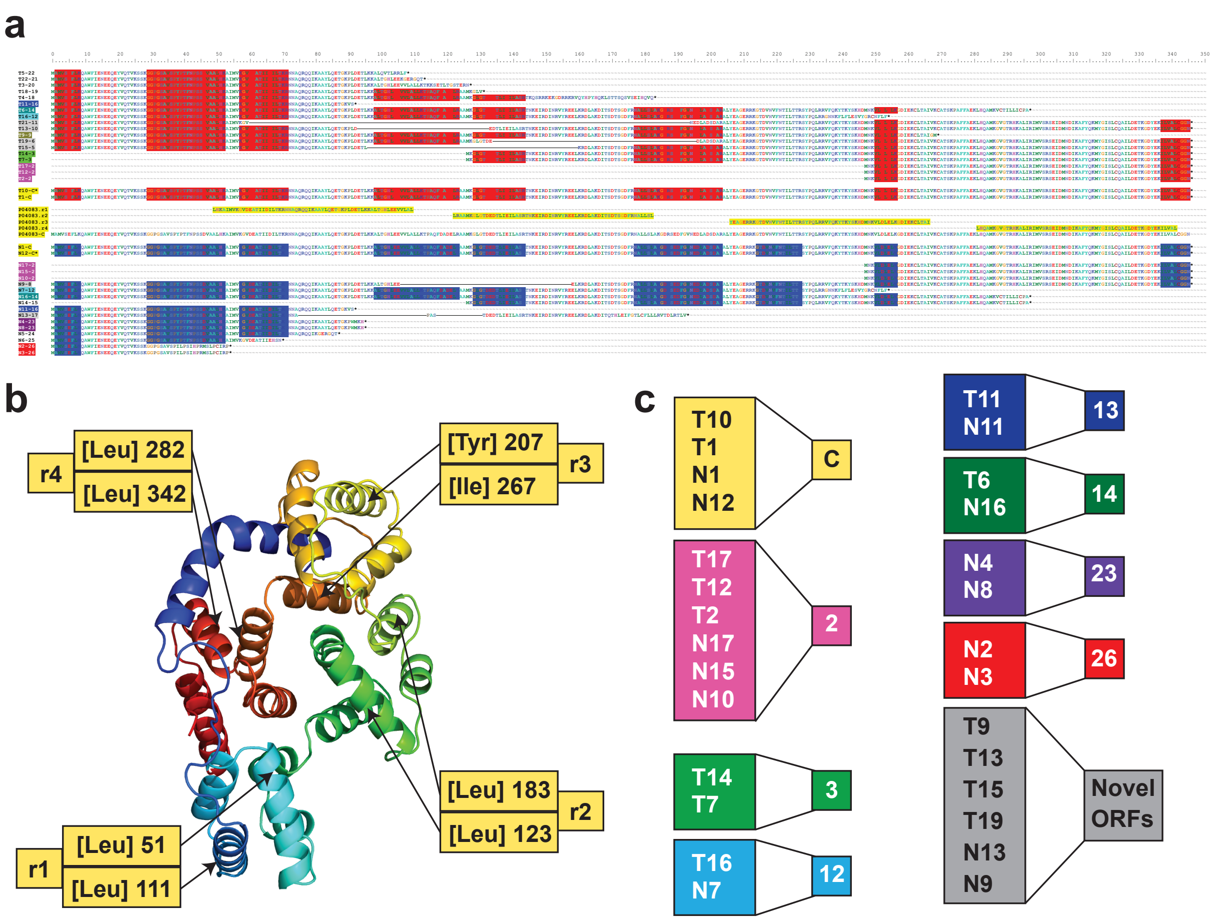


**Figure S1.** Amino acid sequence alignment and proteins predicted from the transcript isoforms identified for *ANXA1* (see **Fig. 4c,d**). (**a**) Amino acid sequence alignments including the isoform identifiers. The canonical protein and conserved repeat domains r1 - r4 are highlighted in yellow. ORFs coding for the same protein are shown in matching colors. Note: Solid lines connecting protein fragments indicate contiguous amino acid sequences predicted from the respective transcript isoform. (**b**) Predicted protein structure of the canonical *ANXA1* protein P04083 with repeat domains r1 to r4 indicated. (**c**) Common predicted proteins for groups of transcript isoforms, ORFs 2, 3, 12, 13, 14, 23, 26 and the canonical ORF (C) are listed and shown with the respective color from panel (**a**). Novel ORFs are indicated.

**(A magnified portion of Figure S1a is on the next page.)**

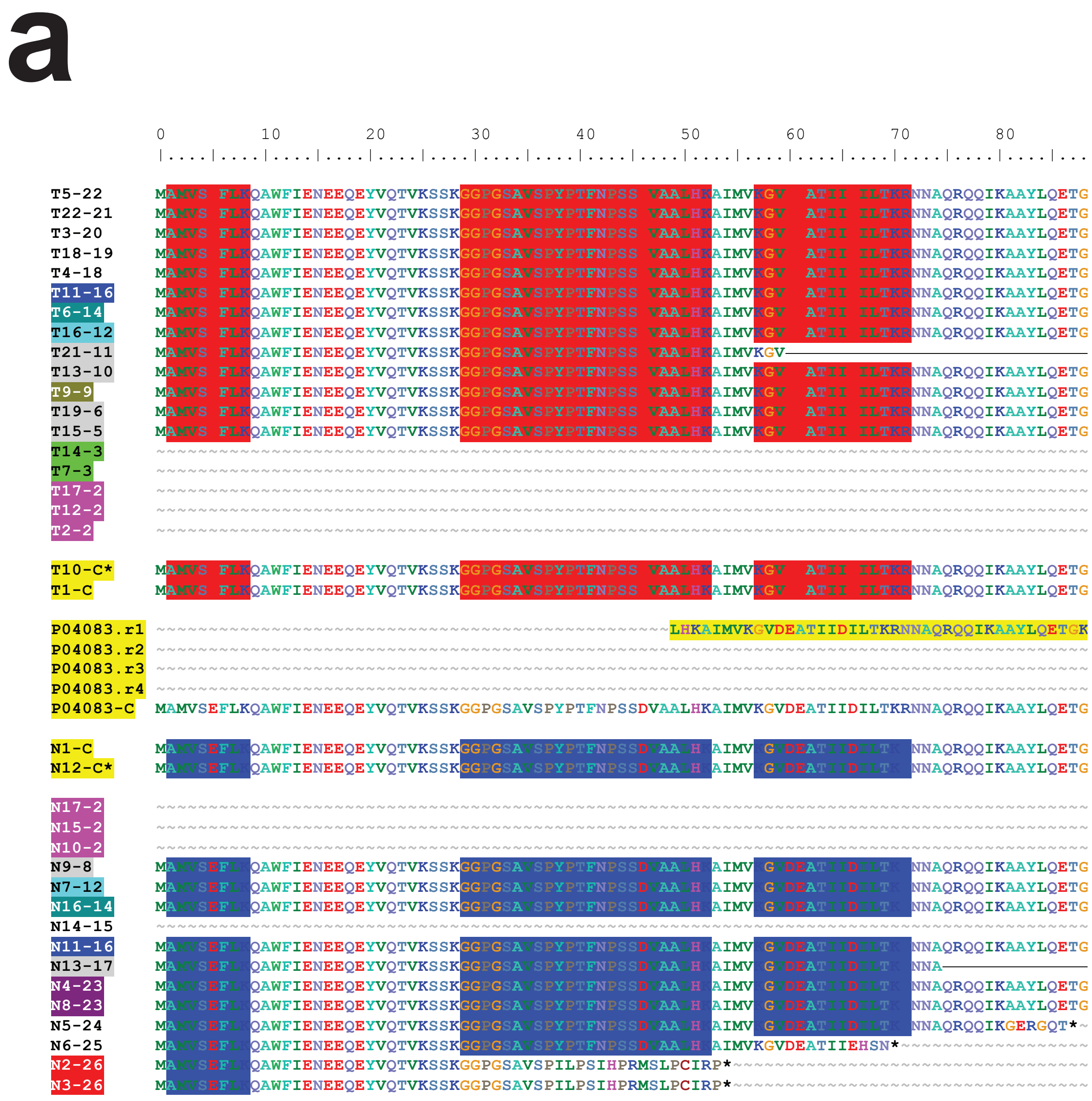

**Figure S1a magnified for better readibility.**

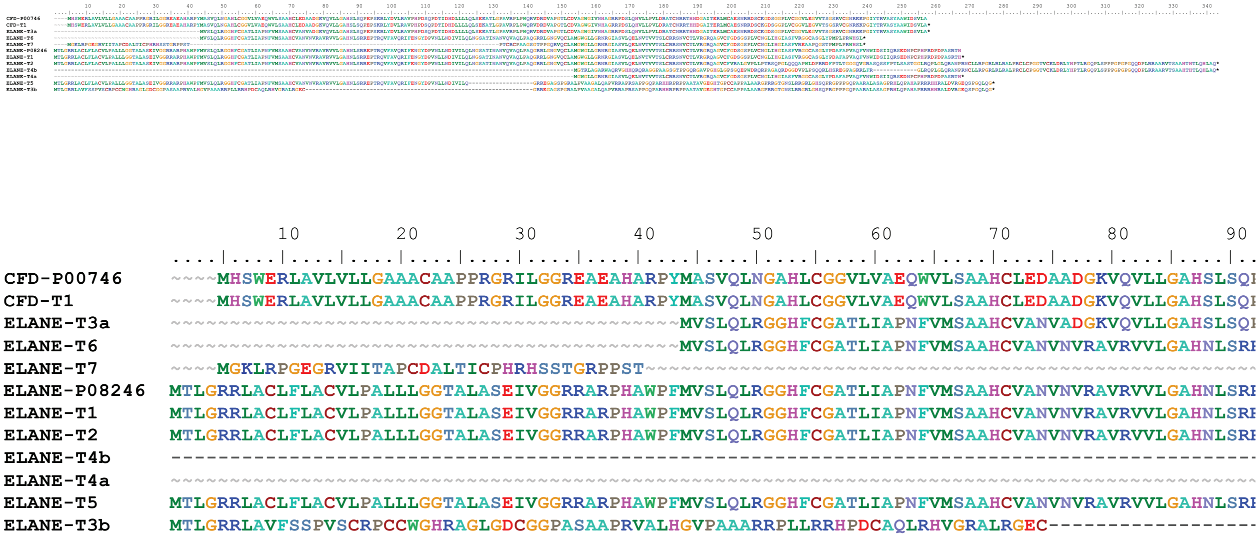


**Figure S2.** Multiple amino acid sequence alignment predicted from transcript isoforms for *ELANE* and *CFD* The alignment is shown in the top panel. The identifiers of the transcript isoforms are included.

**For better readibility a magnified portion of the figure is shown as the bottom panel**.

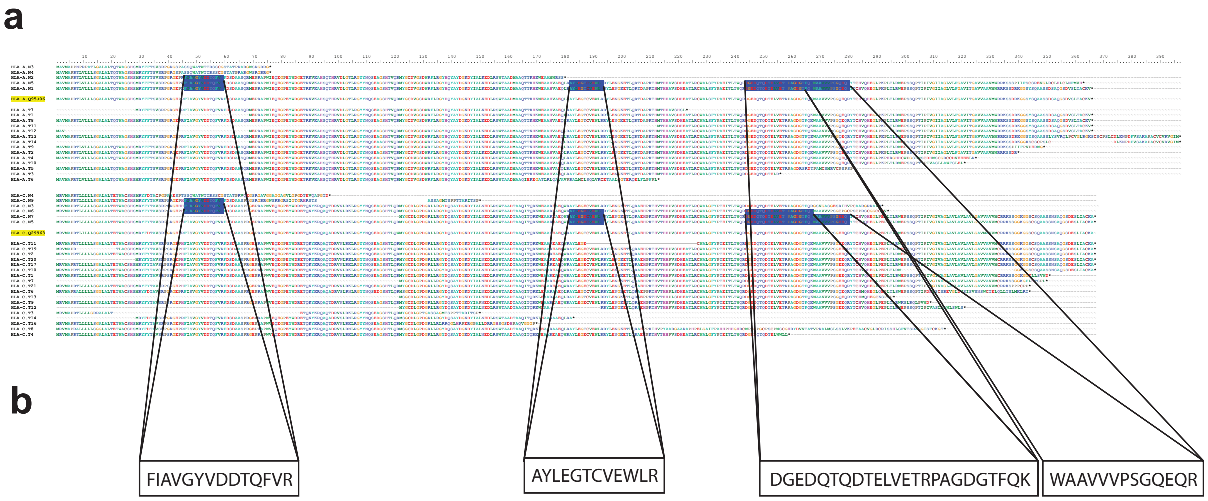


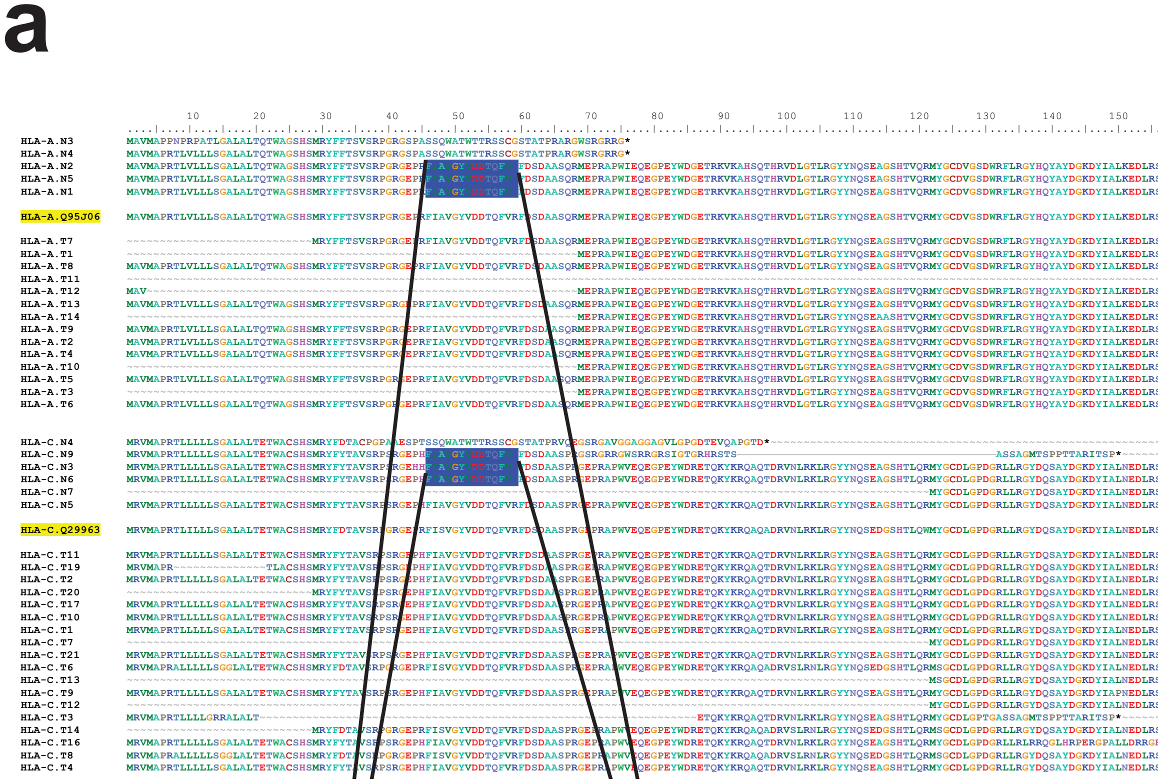


**Figure S3.** Multiple amino acid sequence alignment **(a)** and mass spectrometry detected peptides (**b**) for *HLA-A*, *-B* and *-C* transcripts. **(a)** Sequences predicted from transcript isoforms of *HLA-A, HLA-B* and *HLA-C*. The identifiers of the transcript isoforms are included. Canonical amino acid sequences are highlighted in yellow. **(b)** Peptide fragments identified by mass spectrometry analysis of tryptic fragments of proteins extracted from lin-neg bone marrow cells. The spectra are shown in **Fig. 5d-f.**

**For better readibility a magnified portion of Fig S3a is shown in the bottom panel**

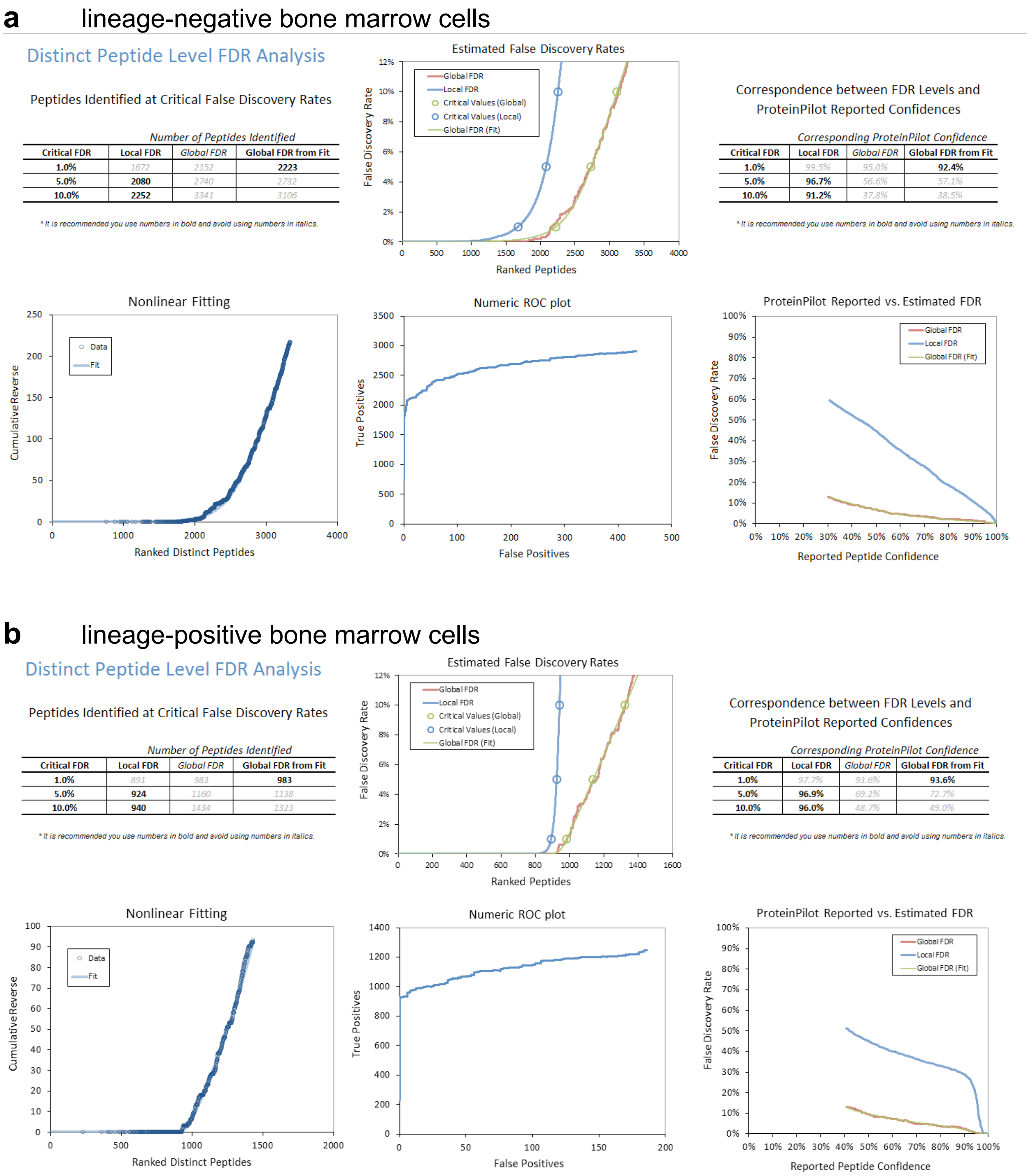


**Figure S4.** Summary of the false discovery rate (FDRs) of the shotgun mass spectrometry of proteins extracted from bone marrow-derived cells. FDR analysis was performed on the ProteinPilot platform (1, 2). The cutoff for positive peptide identification was set at an FDR < 1%. The graphs and tables represent the overview of the proteomics analysis of lineage-negative **(a)** and lineage-positive **(b)** bone marrow cells.
